# Im-seq: Multimodal droplet barcoding enables high-throughput linking of single-cell imaging and gene expression

**DOI:** 10.1101/2025.09.01.672798

**Authors:** Catherine K Xu, Georg Meisl, Nikita Moshkov, Niklas A Schmacke, Karolis Goda, Alexey Shkarin, Maximilian F Schlögel, Sandra Diebold, Christine Schweitzer, Arif Ekici, Tuomas PJ Knowles, Fabian J Theis, Linas Mazutis, Jochen Guck

**Affiliations:** Max Planck Institute for the Science of Light, Erlangen, Germany; Max Planck Zentrum für Physik und Medizin, Erlangen, Germany; Yusuf Hamied Department of Chemistry, University of Cambridge, Cambridge, UK; Institute of Computational Biology, Helmholtz Munich, Munich, Germany; Institute of Biotechnology, Vilnius University, Vilnius, Lithuania; Institute of Human Genetics, Friedrich-Alexander-Universität Erlangen-Nürnberg, Erlangen, Germany; Uniklinikum Erlangen, Erlangen, Germany; Cavendish Laboratory, University of Cambridge, Cambridge, UK; TUM School of Life Sciences Weihenstephan, Technical University of Munich, Munich, Germany; School of Computation, Information and Technology, Technical University of Munich, Munich, Germany

## Abstract

Understanding sequence-phenotype relationships in biology requires large-scale datasets that provide one-to-one mapping of sequence to phenotype. Here, we address this challenge by introducing multimodal droplet barcoding, enabling individual microfluidic droplets to be tracked across both optical and sequencing measurements. We demonstrate the power of this approach through multimodal single-cell analysis. We pair imaging with gene expression profiling of individual cells, identifying genes that correlate with phenotypic features.

## Main text

A central challenge in biology is how function arises from sequence, whether at the level of protein molecules or cells. Fully elucidating this relationship requires large amounts of data linking the modalities of functionality and sequence. This need for detailed data at scale is becoming increasingly urgent as machine learning has progressed rapidly to a level that can now provide unprecedented insights into biological complexity, given the right data [1, 2]. Droplet microfluidics provides an attractive solution where, by compartmentalising reactions in picolitre- to nanolitre-scale droplets, we can achieve orders of magnitude higher throughput than standard well plate-based methods. This has been harnessed extensively in areas including high-throughput single-cell mRNA sequencing (scRNA-seq) and directed evolution of protein functions [3, 4, 5]. However, the true potential of droplets has not yet been realised: connecting multiple single-droplet level measurements of different properties at scale.

Current strategies to link microfluidic measurements across modalities are limited in either information depth or throughput. Sorting droplets into “positive” and “negative” populations as in directed evolution is extremely powerful due to its very high throughput, but reduces optical phenotype measurements to a binary readout [5, 6]. By contrast, static, array-based approaches are inherently limited in throughput and scalability [7, 8, 9, 10, 11, 12]. Similarly, high-precision handling is required in deterministic flow-based systems which precisely control individual cells or droplets, ultimately limiting their analysis rate [13, 14, 15].

Here, we developed *im-seq*, a multimodal droplet barcoding approach as a strategy to address this challenge; we link different data modalities at the single droplet level by tagging each droplet with a physical multimodal barcode (MB) which is identifiable both optically and by sequencing. Hydrogel beads containing DNA barcodes are frequently used to achieve single-cell-level data via sequencing, but these beads are optically indistinguishable [3, 16, 17]. Synthesising a unique, optically distinct bead with a known DNA sequence (an MB bead) for each droplet does not scale favourably. We therefore employ a combinatorial strategy which deliberately harnesses stochasticity, whereby multiple such MB beads are randomly combined in droplets to generate multi-component barcodes [11, 18, 19]; MB beads can be identified in droplets both optically and by sequencing, to yield the complete multimodal barcode (Figure 1a). This combinatorial approach scales extremely well, with just 30 such MB beads able to yield over 1 billion composite barcodes (Figure 1a-d). Here, we used droplet microfluidics to synthesise 41 unique MB bead types from polyacrylamide by varying their size, shape, and colour (Figure 1c, e, f), and reversibly conjugated each type to a unique, known DNA sequence [19, 20, 21].

**Figure 1:**
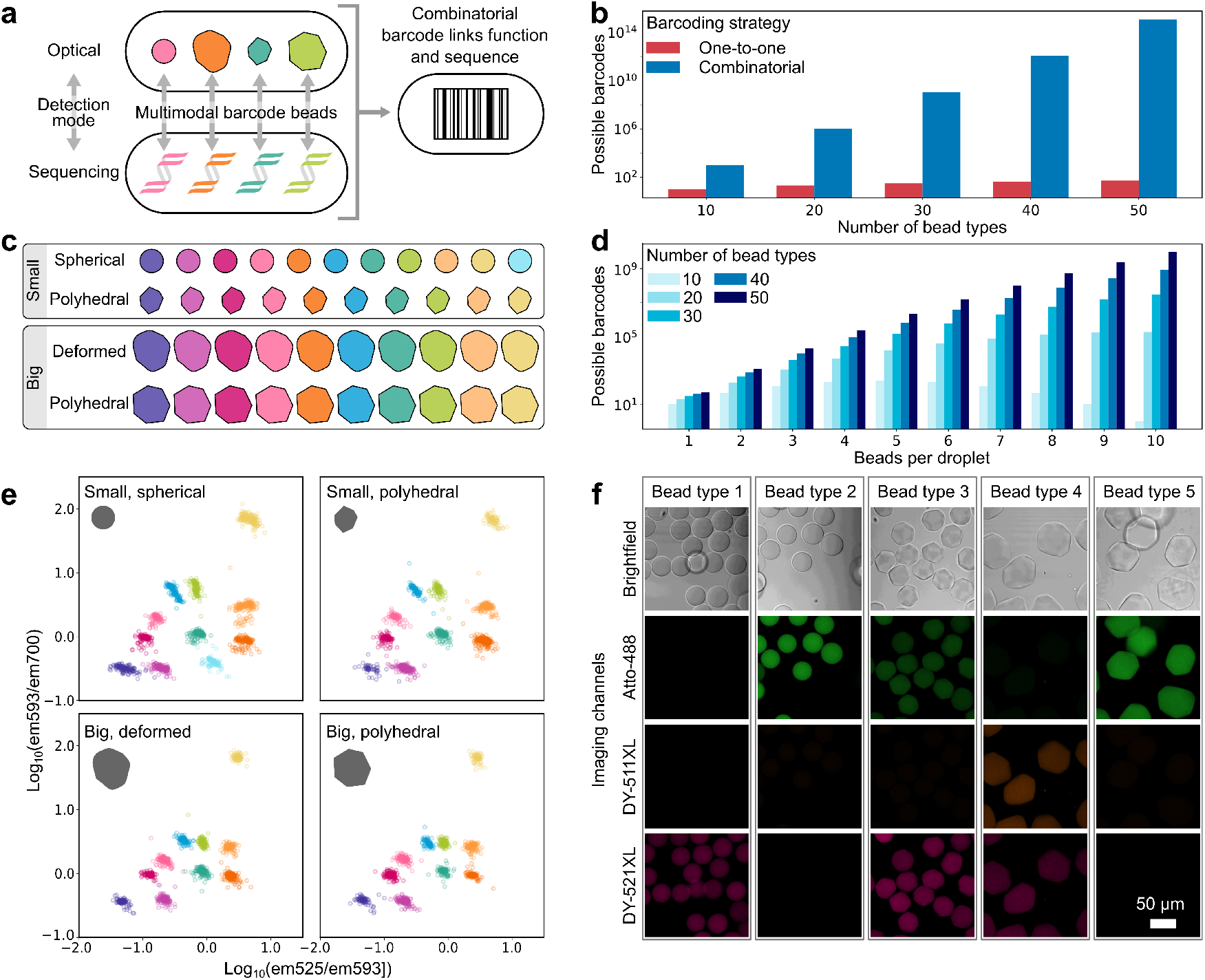
Generation of multimodal barcodes. (**a**) Multimodal barcodes are generated by the random combination of multimodal barcode (MB) beads, each of which is identifiably both optically and by sequencing. (**b**) The number of unique barcodes that can be generated by combinatorics, compared to single bead labelling, shown on a logarithmic scale. Combinations of 30 bead types already produce over 1 billion barcodes. (**c**) Schematic representation of the 41 MB bead types used in this study, using the parameters of size, colour, and shape to render them optically distinct. (**d**) The number of barcodes depends on the number of MB beads per droplet. With 41 MB bead types, millions of barcodes are possible even if the number of beads per droplet is capped at 6. (**e**) Ratiometric bead colours defined by the combination of three fluorescent dyes. The three fluorescence signals (emission at 525, 593, and 700 nm) were reduced to two dimensions by taking the pairwise ratios, removing noise arising from variations in absolute fluorescence intensity. Colours are shown for the four size/shape combinations. (**f**) Example brightfield and fluorescence images of types (columns) of multimodal barcode beads.

Having established the construction of multimodal droplet barcodes, we demon-strated the application of our *im-seq* platform to single-cell analysis. We designed our MB beads to integrate directly into existing scRNA-seq workflows, allowing imaging and transcriptomic measurements to be linked for the same cell. Despite the abundance of technologies for single-cell interrogation, approaches for multimodal characterisation are still sparse. Yet, such datasets are increasingly recognised as crucial for providing both insights into inter-modality relationships and foundations for integrating single-modality datasets. [22, 23, 24, 25]. In contrast to spatial omics methods requiring fixation, our microfluidic characterisation is compatible with live cells and can thus access cell properties such as mechanical stiffness and secretion, and do so in a time-resolved manner [26, 27, 28].

In droplet-based scRNA-seq, single cells are co-encapsulated with uniquely DNA-barcoded beads (blind barcode beads) which index sequences during reverse transcription [3, 4, 16]. Our platform, *im-seq*, images live cells directly prior to their encapsulation in droplets. In addition to blind barcode beads, droplets now also contain MB beads (Figure 2a-c). The blind barcode bead then tags both MB-derived DNA and cell mRNA with the same unique sequence [29, 30]. The combinatorial multimodal barcode thereby links imaging and sequencing data at the single-cell level.

**Figure 2:**
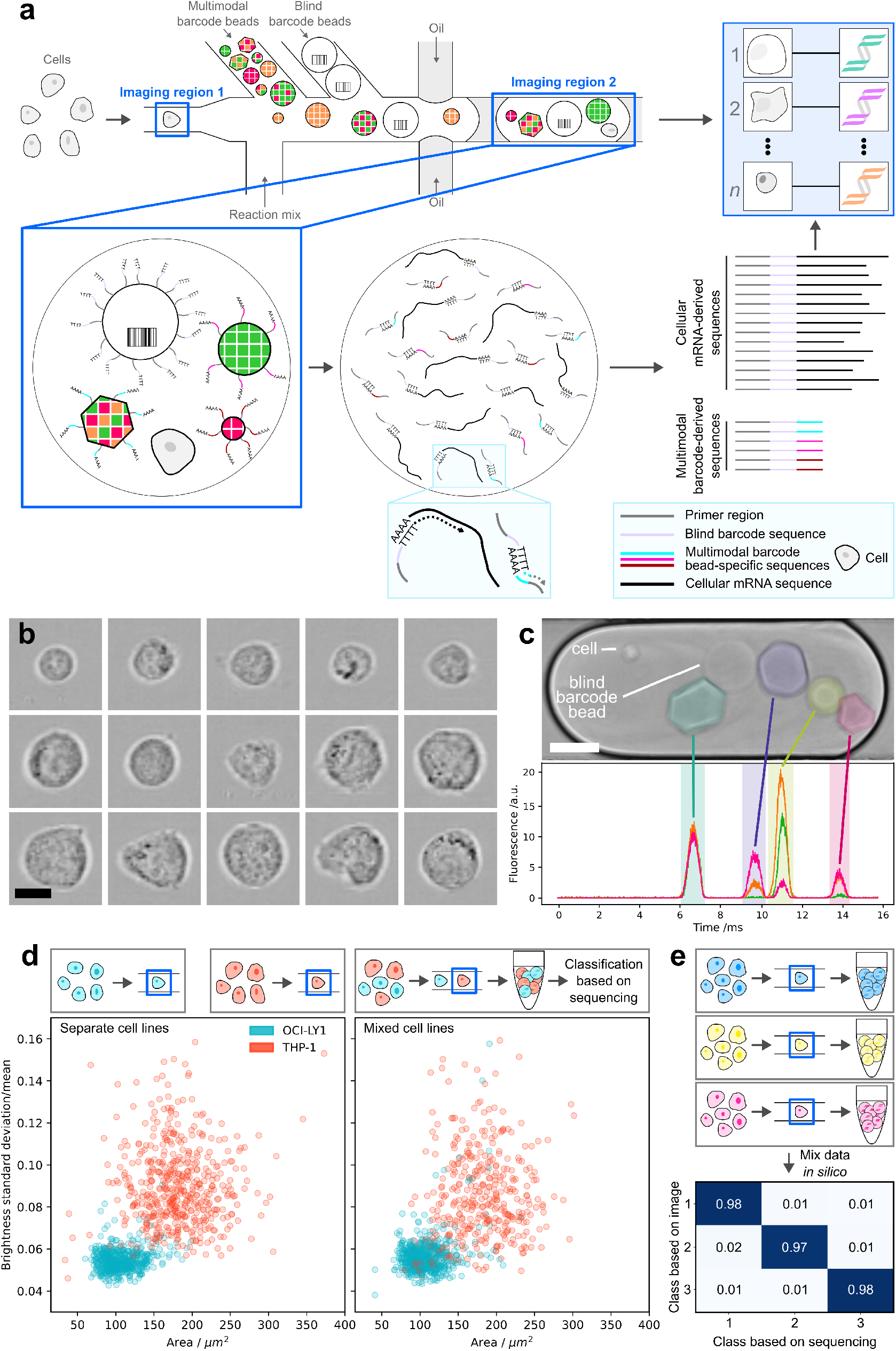
Multimodal barcodes link single-cell image and gene expression. (**a**) Schematic of high-throughput, multimodal single-cell analysis by barcoding. Live cells are imaged (imaging region 1) immediately prior to encapsulation with both blind barcode beads and multimodal barcode beads. The multimodal barcode beads, which are functionalised with known DNA sequences, enable direct image-to-transcriptome linking of single cells. (**b**) Example images of cells (imaging region 1) immediately prior to encapsulation in droplets. Scale bar: 10 µm. (**c**) Droplet containing a cell, blind barcode bead, and four multimodal component beads which comprise the multimodal barcode. Fluorescence traces determine the colour of each bead (coloured as in Figure 1(**e**)) (Scale bar: 50 µm.). (**d**) (Left) OCI-LY1 (turquoise) and THP-1 (orange) cells were imaged in separate measurements, showing distinct physical properties. (Right) OCI-LY1 and THP-1 cells were then mixed together (943 cells in total), and the mixture was measured in a single experiment. The same image features are again shown but points are now coloured according to the gene expression-based cell type classification, validating the image-sequence linking. (**e**) Ground truth validation of linking optical and sequencing data. Data from three separate measurements were mixed *in silico* before linking droplet image and sequence data. In 97% of linked droplets, sequencing data were paired with optical droplet data from the same original measurement, indicating a correct assignment rate of 95% (details in Methods).

To validate that our MB beads correctly link imaging and sequencing data, we applied im-seq to a mixture of two cell lines: OCI-LY1 and THP-1, which differ in physical phenotype at the population level. Labelling the image data according to their linked gene expression profiles, we demonstrated that image and sequence data are linked correctly at the single-cell level (Figure 2d). We performed ground truth experiments to estimate the rate of correct assignment. We pooled data from three independent experiments and performed image–sequence matching without using information about experimental origin. We then compared the resulting assignments with the known experiment labels as ground truth. Over 97% of matched droplets paired imaging and sequencing data from the same experiment, confirming accurate image-sequence matching (Figures 2e, S10), with mismatches likely originating from low signal-to-noise ratios in MB bead sequencing for a minority of droplets.

We then explored gene expression–image feature relationships in OCI-LY1 cells. Compared to the single-modality embeddings of image or gene expression individually (Figures 3a-b, S17, S18), we observe emergent structure in the joint image-gene expression embedding (Figure 3c, S16). For cell area, we see two distinct populations of larger cells emerge, with differences in their morphology (Figure 3c-d). From these datasets, we identified genes whose expression correlated with cell area, which we validated with cell sorting combined with bulk RNA sequencing (Figure 3h-i). Overlaying expression of the two most correlated genes, UBE2C (Figure 3e) and TOP2A (Figure S16), we find that these genes are associated primarily with only one of the two large-area populations, which is only evident with our image-sequence linked data. We investigated additional haematopoietic cell lines, HL-60 and THP-1, with different size distributions compared to OCI-LY1 (Figure 3f-g). While TOP2A was correlated with cell area within each of the cell lines, this association did not extend between cell lines, and we identified additional genes whose image association is driven primarily by differences between cell lines (Figure 3j-l, S13).

**Figure 3:**
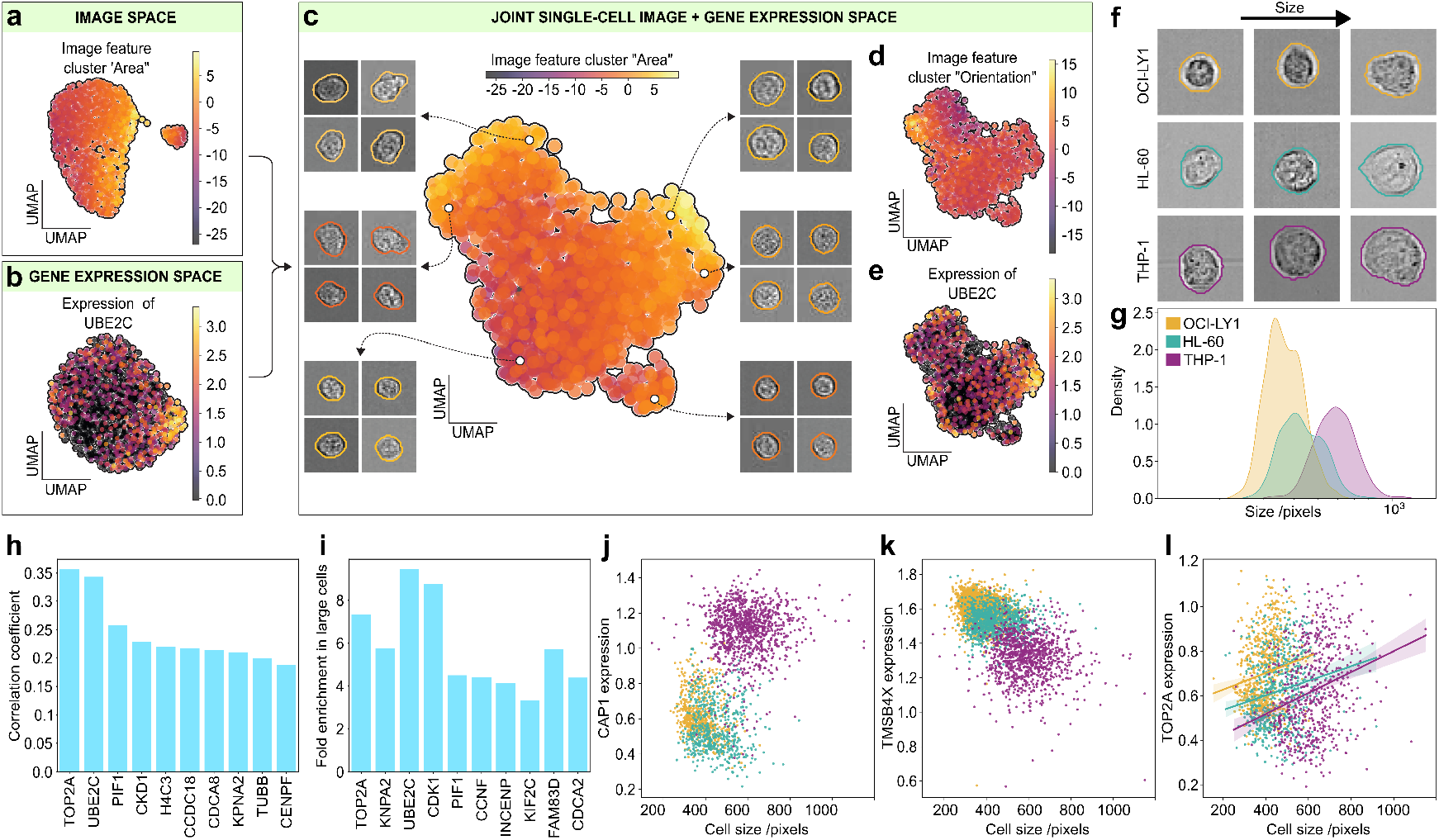
Gene expression-imaging associations in haematopoietic cell lines. (**a**) Principal component analysis (PCA) of single-cell images of OCI-LY1 cells (1652 cells), with points coloured according to the image feature cluster containing cell area. Images were featurised with CellProfiler [31]. (**b**) PCA of single-cell gene expression in the same cells shown in (**a**), with points coloured according to UBE2C expression level. (**c**) Joint single-cell image-gene expression embedding for the same data in (**a**) and (**b**), coloured according to the magnitude of the image feature cluster containing cell area. Example cell images are shown for different regions of the embedding. (**d**) Joint embedding coloured according to the cell image-derived feature cluster containing cell orientation. (**e**) Joint embedding coloured according to relative expression levels of UBE2C. (**f**) Example single-cell images from different cell lines, with each line measured separately, ordered according to cell size and shown alongside their segmentation. (**g**) Size distributions of cell lines as measured by im-seq. Colours throughout the figure are yellow: OCI-LY1, turquoise: HL-60, and purple: THP-1. (**h**) Pearson correlation coefficients for the ten genes most strongly correlated with cell area in OCI-LY1 cells. (**i**) Independent validation of the single-cell findings by bulk RNA sequencing. OCI-LY1 cells were sorted by size, and the largest 5% and smallest 5% populations were analysed by bulk RNA sequencing. Fold enrichment in the largest relative to the smallest cells is shown for the ten most statistically significant genes. Five of these genes also ranked among the ten genes most strongly correlated with cell area in the single-cell analysis (**h**), independently validating the single-cell findings. (**j**-**l**) Examples of different types of correlations between cell size and the expression of individual genes observed by im-seq. Gene expression can be correlated across (**j, k**) or within (**l**) cell lines.

In conclusion, we present *im-seq*, a general strategy for multimodal single-cell characterisation at high throughput. *Im-seq* combines the versatility of image-based characterisation with the scalability of droplet sequencing workflows at the single-cell level (Table S4). Our continuous flow-based approach is fast (currently linked image-sequence data for 1000 cells in 5 minutes) and scalable.

The multimodal barcoding approach is modular and compatible with other imaging flow cytometry modalities beyond brightfield imaging, such as fluorescence imaging [32]. Moreover, the barcodes are compatible with live cells, meaning they can be integrated with functional assays such as time-resolved secretion profiling in droplets [27, 33] and perturbation screens [34, 35, 36], while also avoiding fixation-induced artefacts [26, 37]. We anticipate that multimodal barcoding will open new avenues for studying the relationship between cell phenotypes and gene expression programmes in health and disease, and provide the throughput and information-rich data to power machine learning technologies in biology.

## Acknowledgements

We are grateful to Cornelia Liebers and Manuela Hauke for culturing cells, to Dr Irina Harder, Parth Patel, and Dr Salvatore Girardo for their support in the clean room, and to Dr Rinho Kim for his support with sequencing. We also thank Dr David Morse, Dr Tomasz Kaminski, and Professor Justus Kebschull for their helpful discussions. This research received funding from: the Peter und Traudl Engelhorn Stiftung fellowship (C.K.X), the HFSP Cross-Disciplinary Fellowship (C.K.X.), the Marie Sk-lodowska-Curie Actions fellowship READ-seq (agreement no. 101068803) (C.K.X.). N.A.S. is supported by the Add-on Fellowship of the Joachim Herz Foundation.

## Author contributions

Project conception: C.K.X., G.M., J.G. Experiments: C.K.X., K.G, S.D., C.S. Sequencing: A.E. Software and analysis: C.K.X., G.M., N.M., N.A.S., M.F.S., T.P.J.K., F.J.T. Writing (original draft): C.K.X. Writing (reviewing and editing): C.K.X., G.M., N.A.S., N.M., K.G., A.S., M.F.S., S.D., C.S., A.E., T.P.J.K., F.J.T., L.M., J.G.

