## Supplementary Information for "Im-seq: Multimodal droplet barcoding enables high-throughput linking of single-cell imaging and gene expression"

[†]Deceased

[1]Max Planck Institute for the Science of Light, Erlangen, Germany

[2]Max Planck Zentrum für Physik und Medizin, Erlangen, Germany

[3]Yusuf Hamied Department of Chemistry, University of Cambridge, Cambridge, UK

[4]Institute of Computational Biology, Helmholtz Munich, Munich, Germany

[5]Institute of Biotechnology, Vilnius University, Vilnius, Lithuania

[6]Cavendish Laboratory, University of Cambridge, Cambridge, UK

[7]TUM School of Life Sciences Weihenstephan, Technical University of Munich, Munich, Germany

[8]School of Computation, Information and Technology, Technical University of Munich, Munich, Germany

### Supplementary figures

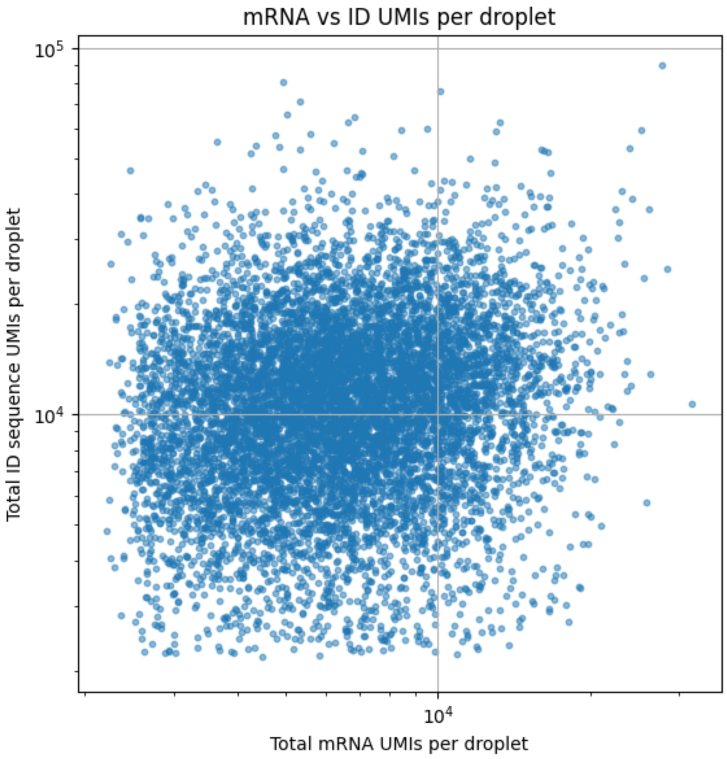


Figure S1: **UMI counts per droplet for mRNA- and optical barcode-derived sequences.** No correlation was evident between UMI counts for mRNA- and optical barcode-derived sequences, indicating that the presence of optical barcode beads does not impact gene expression data by competing for blind barcode bead DNA.

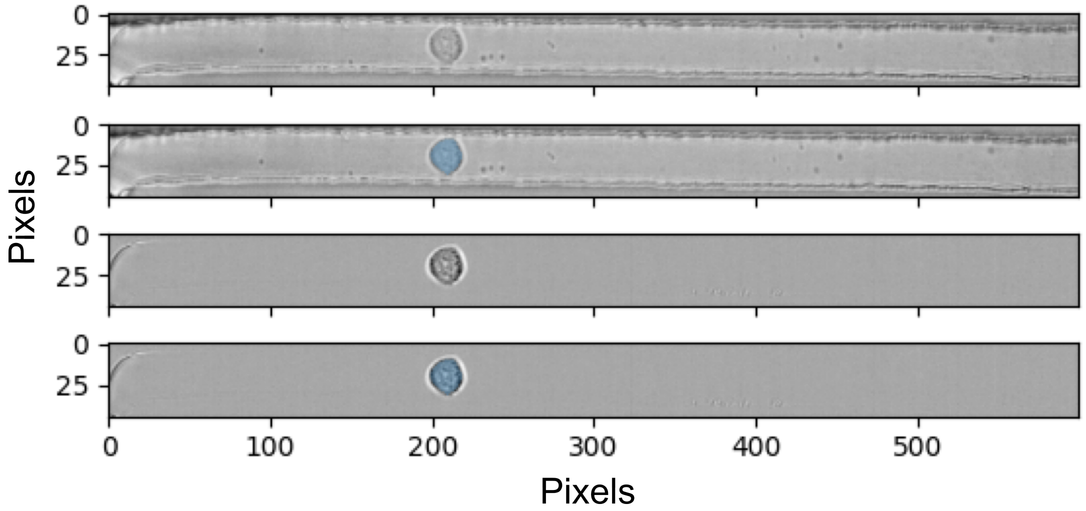


Figure S2: **Background correction and segmentation of cells.** The original image (top two panels) is shown alongside the background-corrected version (bottom two panels), with and without the segmented mask. (Pixel size: 0.68 µm).

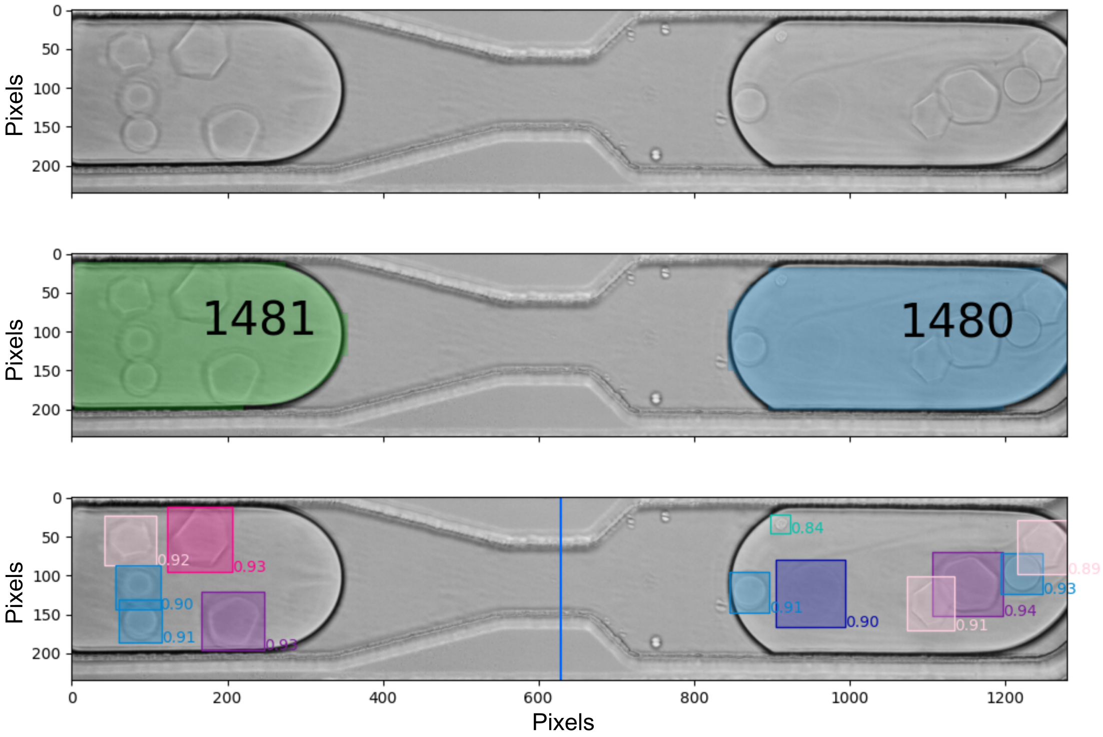


Figure S3: **Image analysis of droplets.** Top: original image. Middle: Segmentation masks for droplets, alongside their assigned id number to be tracked between frames (droplets travel left to right in this image). Bottom: Detection of droplet contents (boxes), shown with the model confidence scores. The object types detected are: cell (turquoise), blind barcode bead (dark blue), and the four size/shapes of optical barcode bead, namely small spherical (blue), small polyhedral (light pink), large polyhedral (dark pink), and large deformed (purple). The blue vertical line indicates the location of the 488 nm laser light sheet.

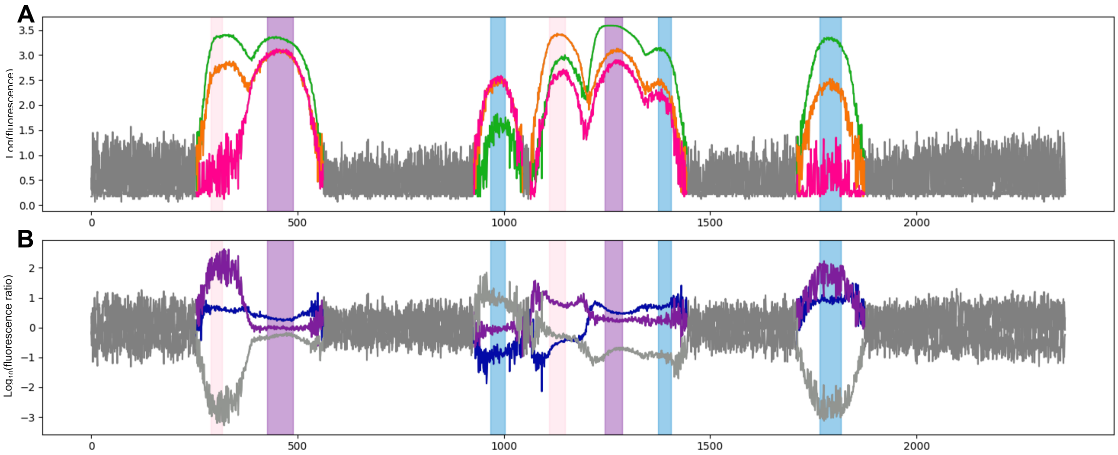


Figure S4: **Determination of optical barcode bead colours.** (**A**) Fluorescence trace for the droplet, with grey regions falling below the signal threshold. Fluorescent regions corresponding to beads were paired with bead types (size and shape) based on their imaged positions relative to the laser light sheet. The individual bead regions detected are highlighted, using the same colour scheme as in Figure S3. Signals are from detectors at 525 nm (green), 593 nm (orange), and 700 nm (pink). (**B**) Pairwise fluorescence ratios: each fluorescence region results in constant ratios to define bead colours (blue: 525 nm/593 nm, purple: 593 nm/700 nm, grey: 700 nm/525 nm).

### SI tables

| Model | Training images | Validation images | YOLO model | Input size | Precision | Recall | mAP@0.5 | mAP@0.5:0.95 | Confidence threshold used during inference |
|---|---|---|---|---|---|---|---|---|---|
| Cell detection | 3756 | 945 | yolov8s.pt | 600x45 | 0.9978 | 0.99221 | 0.995 | 0.85998 | 0.6 |
| Cell segmentation | 3663 | 916 | yolov8n-seg.pt | 640x640 | 0.98945 | 0.99427 | 0.99477 | 0.89921 | 0.7 |
| Droplet segmentation | 954 | 240 | yolov8n-seg.pt | 1280x235 | 0.98778 | 0.98798 | 0.9947 | 0.96565 | 0.2 |
| Bead detection | 2879 | 724 | yolov8n.pt | 1280x1280 | 0.98426 | 0.9904 | 0.99334 | 0.90988 | 0.6 |
| Cell detection in droplets | 23320 | 5834 | yolov8s.pt | 1100x1100 | 0.9916 | 0.98962 | 0.99386 | 0.76073 | 0.5 |

Table S1: Details of ML detection and segmentation models used. Object detection and segmentation were performed using the YOLOv8 architecture (Ultralytics), a deep learning model for instance detection. YOLOv8 was applied in detection or segmentation mode, as indicated. In each case, a custom model was trained on manually annotated brightfield images using the Ultralytics Python framework (Python 3.12.2, PyTorch 2.3.1+cu121). The model was trained for 100 epochs on GPU with a batch size of 16, with data augmentation enabled and optimisation was performed using the default YOLOv8 optimiser configuration. Following inference, objects were filtered according to the given confidence thresholds.
